# Larval thermosensitivity shapes adult population dynamics in *Anopheles* mosquitoes

**DOI:** 10.1101/2023.09.19.558414

**Authors:** Juan Estupiñán, Anna M. Weyrich, Paula Schlösser, Charlene Naujoks, Markus Gilden-hard, Assetou Diarra, Mouctar Diallo, Djibril Sangare, Arndt Telschow, Chih-hao Hsieh, Elena A. Levashina, Paola Carrillo-Bustamante

## Abstract

Mosquitoes are vectors of human life-threatening pathogens, posing a significant global health threat. While the influence of temperature on mosquito life-history traits has been extensively studied in laboratory settings, the ecological factors shaping mosquito development and population dynamics in natural environments remain poorly understood. Here, we used a multi-disciplinary approach, integrating field data from Mali, laboratory experiments, and mathematical modeling, to investigate the causal relationships between climate variables and the abundance of *Anopheles gambiae s*.*l*. mosquitoes. Using convergent-cross mapping analyses an adult abundance in the Nanguilabou village, we observed that the dynamics of adult mosquito populations was driven by larval thermosensitivity. To elucidate the underlying mechanisms, we conducted experimental studies that revealed a density-dependent larval thermal response. Through mathematical modeling, we quantified the complex interplay between temperature and larval density, demonstrating that temperature and density have independent, non-synergistic effects on larval developmental speed, mortality, and pupation rates. Our findings provide a mechanistic understanding of how larval development shapes adult mosquito populations, highlighting the significance of multidisciplinary approaches in studying climate-driven mosquito population dynamics.

## Introduction

Vector-borne diseases constitute a major public health concern worldwide, accounting for 17% of all infectious diseases and causing 700,000 deaths annually (World Health Organization 2020). Mosquitoes, the vectors of life-threatening pathogens including dengue virus and malaria parasites, exhibit a complex life cycle consisting of aquatic (juvenile) and terrestrial (adult) stages. Consequently, mosquitoes are exposed to a wide range of biotic (inter- and intra-specific competition, microbiome) and abiotic factors (temperature, rainfall, humidity) and are highly sensitive to environmental changes in their habitats (Deutsch *et al*. 2008; Mordecai *et al*. 2019). Understanding the mechanisms through which the environment affects each mosquito’s life cycle stage and regulates population dynamics is critical to predicting mosquito-borne transmission during future climate change scenarios.

Mosquitoes are ectotherms and therefore respond to changes in temperature by modulating their basic life-history traits, including development, survival, fertility, and biting rate (Delatte *et al*. 2009; Deutsch *et al*. 2008; Franklinos *et al*. 2019; Huxley *et al*. 2022; Mordecai *et al*. 2019; Paaijmans *et al*. 2013). While the thermal response of mosquitoes has been well characterized, the effects of other ecological variables on mosquito life-history traits have been only scarcely investigated (Brown *et al*. 2023; Huxley *et al*. 2021; Huxley *et al*. 2022). Moreover, mosquitoes’ thermal biology is typically studied in controlled-laboratory conditions. To which extent these findings can be generalized to natural field conditions remains unknown (Deutsch *et al*. 2008; Metcalf *et al*. 2017).

Quantifying the environmental effects on mosquito life-traits and population dynamics in natural population represents a challenge. Multiple studies have used time series collections of mosquito abundance across geographical locations linking mosquito population dynamics to climate variables including humidity (Alencar *et al*. 2015; Bashar *et al*. 2014; Santos-Vega *et al*. 2022), rainfall (Barrera *et al*. 2011; Bashar *et al*. 2014; Chaves *et al*. 2012), and temperature (Alencar *et al*. 2015; Barrera *et al*. 2011; Chaves *et al*. 2012; Scott *et al*. 2000). However, the commonly used statistical time-series analysis can only provide associations based on linear correlations (Metcalf *et al*. 2017). As correlation does not necessarily represent causation, it is unclear whether changes in these variables would indeed cause shifts in mosquito abundance.

The complex, non-linear nature of mosquito-environment interactions calls for new experimental and computational methods to characterize how climate factors affect mosquito life-history traits. Here, we use a multi-disciplinary approach that combines time-series data of daily abundance of malaria vectors *Anopheles gambiae s*.*l*. in the field, laboratory data, and mathematical modeling. The combination of field and laboratory studies: (i) detected causal relationships between mosquito abundance and two environmental variables: temperature and dew point; (ii) identified the mechanisms by which temperature and larval density regulate mosquito development. Our time-series analysis revealed a delayed causal effect of water temperature and dew point on mosquito population dynamics, suggesting a stronger environmental footprint on immature stages than on adult mosquitoes in the field. Additionally, our experimental and modeling results identified the role of temperature in larval death rate and duration of larval development. Interestingly, we found that larval thermosensitivity was modulated by density. Taken together, our findings suggest that the larval life history traits are shaped by individual effects of environmental forces and propose a mechanistic understanding of how the environment may drive the abundance of mosquito populations in the field.

## Results

### Water temperature and dew point drive mosquito abundance in natural populations

To study environmental effects on mosquito populations, we used time-series data of adult mosquito abundance (*Anopheles gambiae s*.*l*.) and five weather variables (air and water temperature, rainfall, relative humidity, and dew point) collected in Nanguilabougou, Mali, during the 2015 rainy season (Gildenhard *et al*. 2019) (fig.1a). We explored the causal effects of the weather variables on *Anopheles* population dynamics with convergent cross-mapping (CCM), an empirical dynamic modeling approach that makes minimal assumptions about the underlying biological mechanisms of the system (Sugihara *et al*. 2012). This method assumes that if a time series (*x*) influences another (*y*), then values of *x* are encoded into *y* and can therefore be reconstructed from *y* alone. Thus, the causal effect of *x* on *y* is determined by how well *y cross-maps x*. Causality is detected if the cross-mapping skill (*ρ*) increases monotonously with the amount of data used, converges to a maximal value *ρ*_*max*_, and *ρ*_*max*_ is significantly higher than that of a null (random) model (fig. 1b) (Sugihara *et al*. 2012). See Methods for a detailed description.

**Figure 1:**
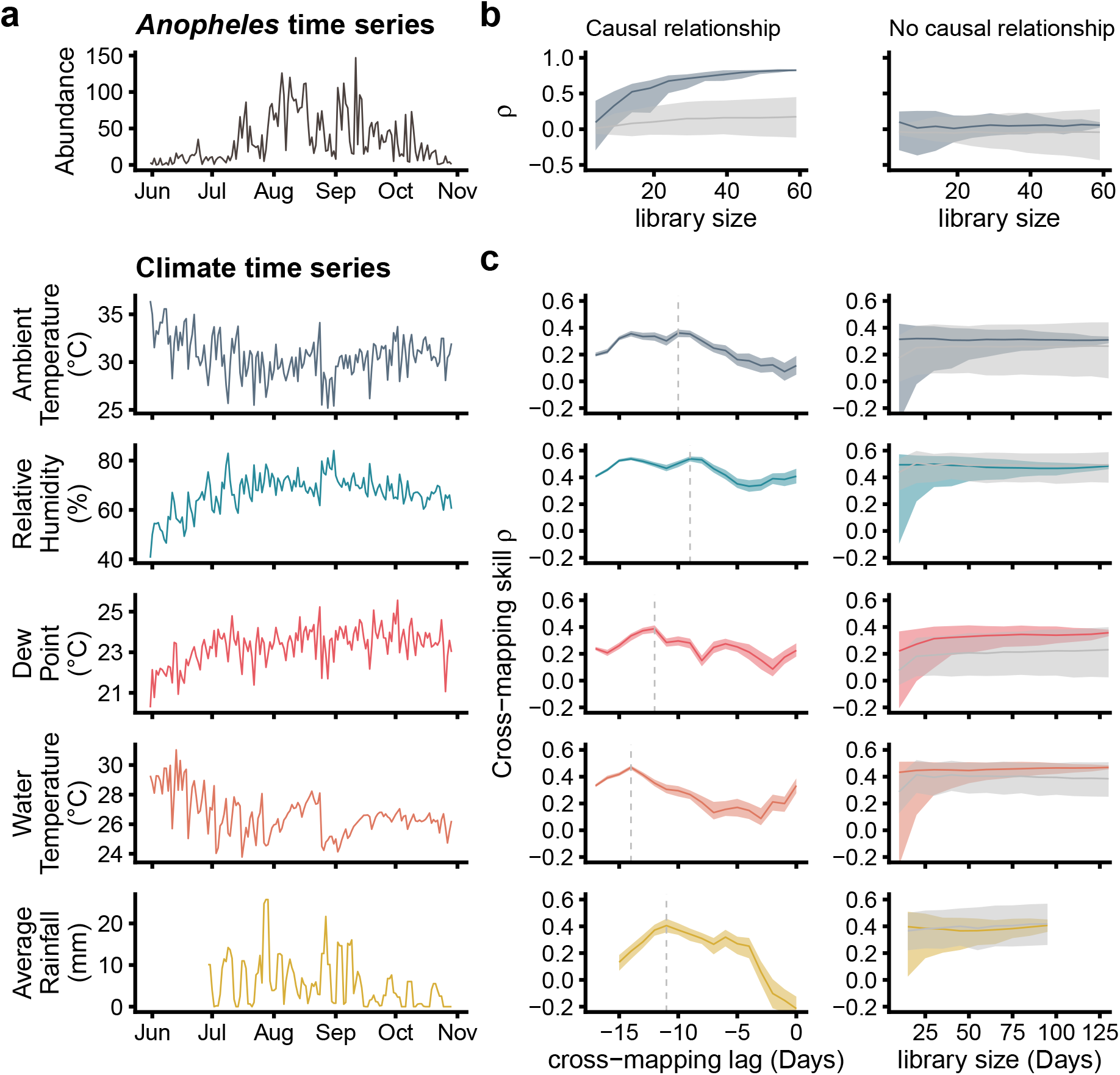
Water temperature and dew point drive *Anopheles gambiae s*.*l*. abundance in natural populations.**(a)** Dynamics of mosquito abundance, ambient temperature, relative humidity, dew point, water temperature, and rainfall in Nankilabougou, Mali, during the 2015 rainy season. Mosquitoes and mean climate variables (3-day rolling average for rainfall) were collected daily. The rainfall time series is shorter due to data loss. **(b)** Causal relationship between the environment and mosquito populations were tested with empirical dynamical modeling. In the presence of causal relationships, the cross-mapping skill *ρ* increases and converges to a maximum value when increasing the time series. *ρ* is considered significant if it is higher than the skill of a null model (light gray regions). **(c)** Delayed causal relationships were analyzed by lagged cross-mapping of the environmental variables between 0 and 17 days. *ρ* converged and was significant only for the dew point and water temperature time series. Solid lines represent the median out of 500 bootstrapped cross-mapping reconstructions, shaded areas indicate the 2.5 and 97.5 percentiles.

We found no immediate causal effect of any environmental variable on the abundance of adult mosquitoes (fig. S2). Instead, the cross-mapping skill increased when applying a time lag to each climate variable (fig. 1c), indicating that historical values of the environmental variables were encoded in the adult mosquito data. Only dew point and water temperature showed a significant causal effect with a delay of 12 and 14 days, respectively. As a control for spurious relationships, we tested nonsensical directions of causality (mosquitoes influencing climate) and none showed significance (fig. S3). Because the delays of water temperature and dew point correlate with the expected duration of mosquito development from an egg to an adult (10-14 days), our CCM analysis suggest that the environment has a stronger effect on larval stages than on adult mosquitoes.

### Combined effects of temperature and larval density determine the dynamics of larval development

CCM can detect causal relationships between two variables, however, it can neither assess the strength nor the direction (positive or negative) of these interactions. To study how the environment affects larval development, we designed an experimental setup that assesses the transition from larvae to adults under a series of temperature regimes. Previous studies have indicated that larval density may also impact the abundance of adult mosquito populations (Huxley *et al*. 2021; Huxley *et al*. 2022). Given the fluctuating nature of larval density in the field breeding site, we explored the combined effects of temperature regimes and larval densities on adult emergence. Specifically, we reared larvae at five temperatures (24, 26, 28, 30 and 32°C) and three densities (100, 250, and 500 individuals per 700 mL of water, fig. 2a). Due to the large number of larvae used in the experiments, it was not feasible to track individual larvae during the early developmental stages (L1-L3). Consequently, the transitions between instar stages and death events were not measured explicitly. We quantified the final steps of larval development by counting daily the number of pupae until the last remaining larva developed or died (fig. 2b), assessing the day of pupation start, the duration of pupation from the start day to the last emerged pupae (pupation span), and the overall percentage of emerged pupae (fig. 2c-e).

**Figure 2:**
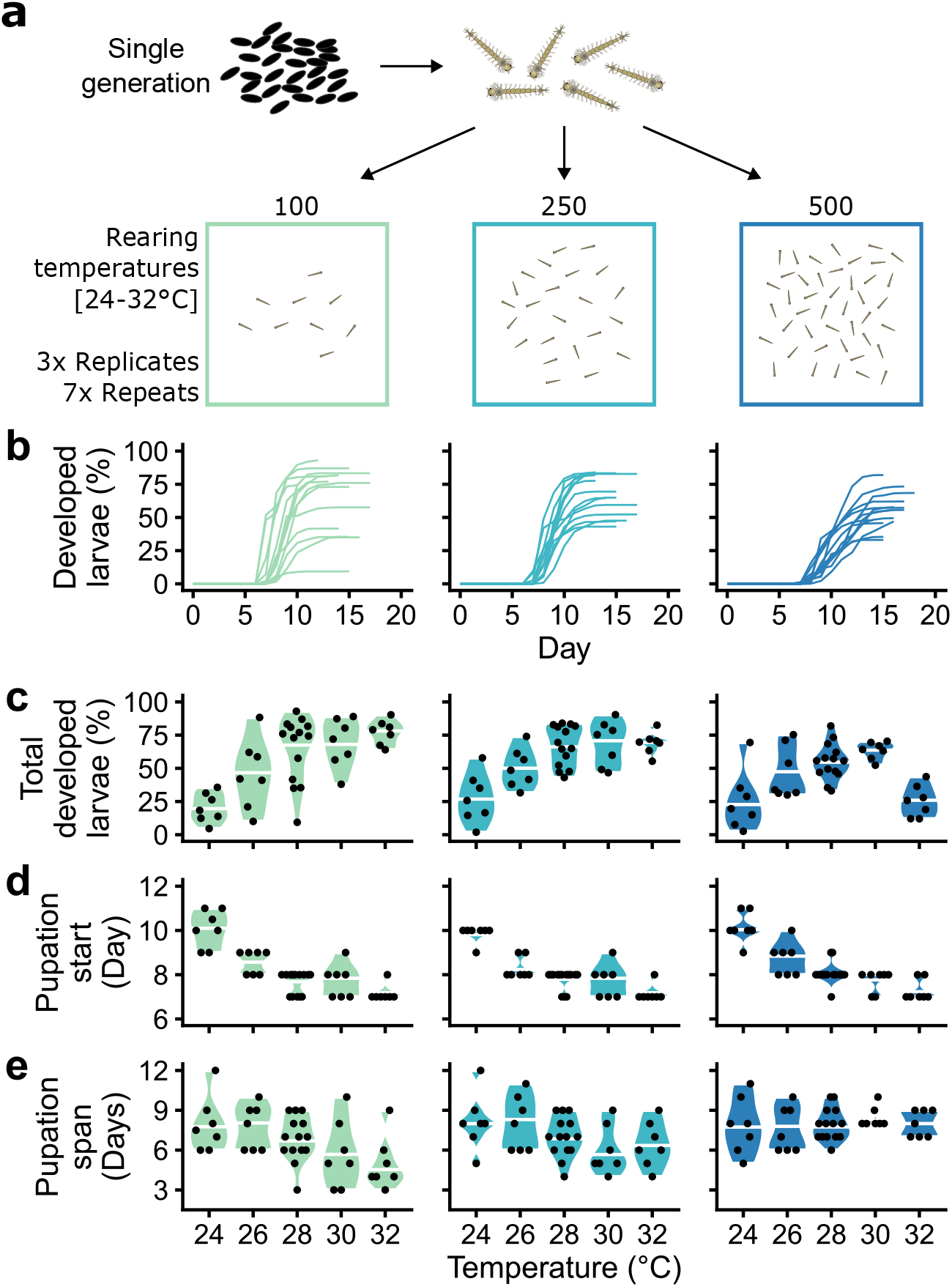
Temperature and larval rearing density determine larval development. **(a)** Experimental design scheme. Larvae derived from one single generation were reared in parallel at three densities: 100 (cyan), 250 (blue), and 500 (dark blue) larvae per 700 mL of water; and five temperatures: 24, 26, 28, 30, and 32°C. To study the effects of these 5 temperature values, experiments were carried out in two groups: 24, 26, 28°C, and 28,30, and 32°C. Each experimental condition was done in triplicates and repeated seven times (n=3, N=7). **(b)** Representative time series of the cumulative developed pupae from T=28°C. Analysis of each development curve provided **(c)** the final percentage of larvae that developed into pupae, **(d)** the start of pupation, and **(e)** the pupation span, i.e., the duration in days from the start to the last day of pupation. For all plots, each dot represents the median of three technical replicates. Violin plots represent the range and median (horizontal white line) of the experimental repeats.

We observed that larval development exhibits a non-linear thermal response, where the optimal rearing temperatures were dependent on larval density. Specifically, the larvae reared at low densities (100) displayed optimal growth at high temperatures (32°C), whereas high densities (500) shifted the optimal temperature to 30°C. Furthermore, increasing temperature accelerated larval development by shifting the start of pupation from 10 days at 24°C to 7 days at 32°C, irrespective of larval density. In addition, higher temperatures shortened the pupation span only at low densities (100 and 250). Overall, our experimental findings suggested that temperature effects on larval development and pupation span were modulated by density, but the exact physiological mechanisms and contribution of each variable remained unresolved.

### Temperature and density affect different traits in larval development

To investigate how temperature and density mechanistically affect larval development we used an unbiased computational analysis. We constructed a mathematical model describing a population of larvae (*L*) that die at a constant rate *δ* and develop into pupae (*P*) at a rate *γ*, after a time *t*_*on*_. Allowing every parameter to change depending on temperature, density, or a combination of both, we obtained 16 different models of environment-dependent larval growth (see eq.1 in Methods). All models were ranked according to their performance to fit the experimental data (table S1).

The best-performing model predicted that larval growth and pupation were influenced by density and temperature, while larval mortality was exclusively thermosensitive (table 1). These results suggested that high-density rearing conditions delayed larval development without affecting larval mortality. The model showed a high level of agreement between the observed data and predicted outcomes (fig.S5a), and was further validated through an out-of-sample dataset encompassing two densities (20 and 1,000 larvae per pan at 28°C, fig.S5b). Notably, none of the models in which temperature and density co-modulated the same parameter produced accurate predictions, implying that these two environmental factors independently affected the examined larval traits. In summary, our results showed that optimal larval development is determined by mortality and developmental rates, which are modulated by the non-synergistic effects of temperature and larval density.

**Table 1:**
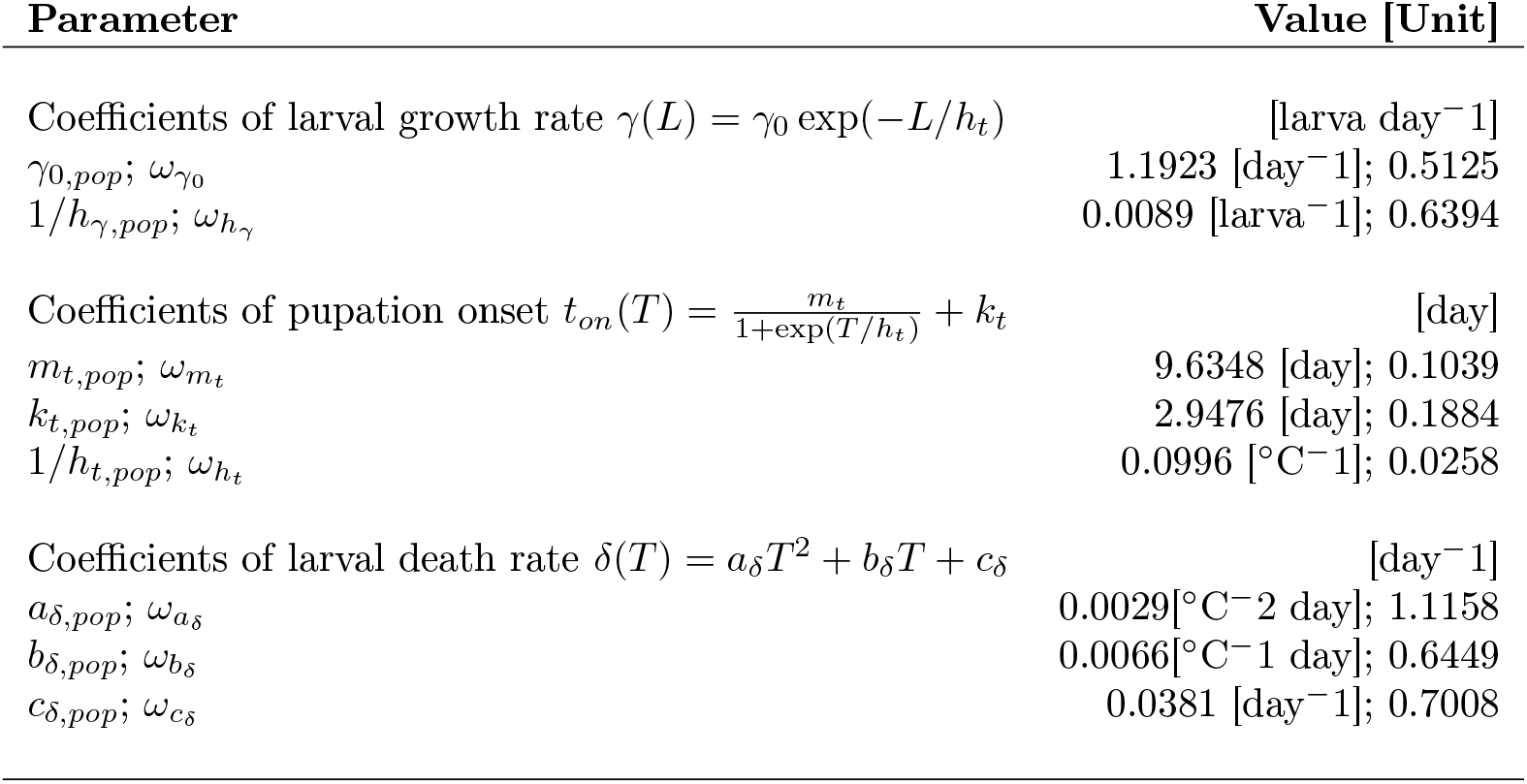
Parameter estimates for the best-performing larval development model. All parameters were estimated using non-linear mixed-effect models. Here, *μ*_pop,*i*_ represents the population estimate for each parameter and exp(*ω*_*i*_) the individual deviation from the population. See Methods for a detailed description.

## Discussion

To elucidate the mechanisms that drive mosquito populations, we used a multidisciplinary approach that integrated field time-series data collections, laboratory experiments, and mathematical modeling. Our combined results suggested that *Anopheles gambiae s*.*l*. population dynamics is shaped by water temperature, dew point, and larval density indirectly through modulation of larval developmental speed, mortality, and pupation rates.

We used CCM to extract the environmental drivers from field-collected data, a method which has been successfully applied in recent ecological and epidemiological studies (Deyle *et al*. 2016; Grziwotz *et al*. 2018; Nova *et al*. 2021). Our CCM analysis was based on daily measurements collected during a single rainy season, providing insights into the dynamics at this specific time resolution. One limitation of our study is that it was based on dense time-series data from only one season. Therefore, it is possible that other factors that were not detected here impact adult mosquito abundance at longer timescales (Jian *et al*. 2016). Therefore, larger datasets that span several seasons will be needed to capture long-term variations. Our findings underscore the necessity for better-designed field studies encompassing daily sampling over multiple seasons to fully exploit the potential of CCM, thereby enabling a more comprehensive understanding of the environmental drivers of mosquito abundance.

Importantly, our findings from CCM revealed the essential need to consider the dynamics of larval populations which is difficult to quantify in the field. Here, we complemented our field data on adult abundance with controlled laboratory experiments to reveal the mechanisms by which the environmental factors impact larval life-history traits. Although measuring individual stage-specific development and survival rates in these experimental settings was not possible, we overcame this limitation by using mathematical models that extract and quantify mechanisms caused by temperature and density changes. Through the construction of multiple models and their fitting to the experimental data, we tested multiple hypotheses and gained unique insights into the dynamics of larval populations. We found that temperature exhibited a stronger effect on larval population dynamics as compared to density. In agreement with previous studies (Bayoh *et al*. 2004; Christiansen-Jucht *et al*. 2015; Kirby *et al*. 2009), our results showed that higher temperatures shortened larval developmental rates and increased larval mortality. The combination of these effects resulted in a unimodal thermal response with an optimum observed at intermediate temperatures. Notably, the non-linear response was modulated by larval density, as also observed for *Aedes* mosquitoes (Huxley *et al*. 2021; Huxley *et al*. 2022). Importantly, our model showed that the two variables, namely temperature and density, acted independently and generated a non-linear response of the studied larval traits.

When examining the effects of density on larval traits, we were surprised to find out that higher densities delay larval growth rates but do not cause mortality in laboratory conditions. The model predictions were validated through additional experiments using variable larval densities, confirming that even at high densities larval developmental delay was not linked to mortality. These findings challenge the existing assumptions typically used in mathematical models that either ignore density-dependence (Agusto *et al*. 2017; Arifin *et al*. 2014; Eckhoff 2011; Ewing *et al*. 2016; Reiner *et al*. 2013) integrate it in a simplified manner (Reiner *et al*. 2013), or assume density-dependent mortality (Depinay *et al*. 2004; Natiello *et al*. 2020). In contrast, our study demonstrated that major density-dependent effects modulate larval developmental rates and thermal responses. Mechanistic understanding of the environmental effects on mosquito life-cycle traits is important to avoid wrong assumptions that may hinder the ability of a mathematical model to capture population dynamics (Cator *et al*. 2020; Walker *et al*. 2021). Therefore, our findings reported here together with the availability of high temporal resolution data sets should benefit future mathematical models that explore functional responses of mosquito life-history traits.

We did not explicitly explore species-specific responses of sympatric mosquito populations to environment. However, our findings on thermosensibility of *Anopheles* mosquitoes differ from the earlier studies that reported the higher density-dependent sensitivity of *Aedes* larvae as compared to temperature (Hancock *et al*. 2016). These observations highlight that many assumptions cannot be simply generalized across all mosquito species.

Our study demonstrated the critical role of larval thermosensitivity in regulating *Anopheles gambiae s*.*l*. abundance. This finding is in line with our previous report that did not find evidence that temperature, humidity, or rainfall directly shape the abundance of adult mosquito populations (Gildenhard *et al*. 2019) and with recent studies in Burkina Faso that uncovered a correlation between rainfall and temperature with *Anopheles gambiae s*.*l*. abundance with a time delay of 1-3 weeks (Taconet *et al*. 2021). All these results taken together, strongly support the importance of immature stages in driving mosquito abundance in the field and demonstrate the power of multi-disciplinary approaches. Importantly, the newly-generated knowledge on the mechanisms that shape population dynamics in the field should provide a valuable resource for epidemiological modelling to predict how vector-borne diseases may evolve under future climate change scenarios.

## Material and methods

### Field data collection

The description for the time-series collection is fully explained in a previous publication (Gildenhard *et al*. 2019). Briefly, mosquito collections were performed in Nankilabougou, Mali, during the rainy season (June -November) of 2015. In total, 5,171 *Anopheles* mosquitoes were collected in a period of 152 days using CDC light traps. From them, 4,200 were identified as *A. coluzzii*, 516 as *A. gambiae*, and 455 as other/unidentified. Ambient temperature, humidity, and dew point data were obtained by a data logger placed in the village, water temperature was measured from the main water body near the village. Rainfall data were obtained from the Bankumana weather station located around 6 km from Nankilabougou. For the rainfall time series, a rolling average of 3 days was applied representing accumulated water in larval habitats.

### Larval development in experimental setting

Freshly hatched first stage *Anopheles coluzzii* (*Ngousso* strain) larvae (L1) were reared in plastic containers (75 x 250 x 250 mm) at variable temperature (24, 26, 28, 30, and 32°C) and density (100, 250, and 500 larvae per 700 mL demineralized water) in plastic containers (75 x 250 x 250 mm). Larvae were fed daily with the 2.5 mL of the mix (2:1:1 ratio) of Bovine liver powder (NOW foods, USA), Tuna meal (Common Baits GmbH, Germany), and VanderzantTM Vitamin Mix (Sigma-Aldrich, Germany). Larval development was monitored daily by counting the number of larvae that converted into pupae. The measurements stopped when the last remaining larvae pupated or died. Experiments were carried out in two groups: 24, 26, 28°C, and 28,30, and 32°C. We used 28°C as the control in every group as it is the typical rearing temperature in our insectary. Each experimental condition was done in triplicates and repeated seven times (n=3, N=7).

### Mathematical Analyses

#### Convergent cross mapping (CCM)

We used convergent cross-mapping (CCM) for determining the existence of causal forcing from the climate variables to the mosquito abundance. In short, the dynamical properties of a complex system can be reconstructed by embedding individual system variables (for example, *x* or *y*). If there is a correspondence between the historical values of variable *y* and the state of *x*, then it can be inferred that the information of *x* is encoded into the *y* time series, and thus, *x* is a causal variable to *y*. Within the CCM framework, we determined the embedding dimension (fig. S1a) and the degree of non-linearity (fig. S1b) for each of the variables before proceeding to cross-map the mosquito abundance manifold to each climate variable. To consider the potential delayed response of mosquito abundance to the forcing climate variable, we tried the lagged effect up to 17 days (15 days for the rainfall time series). We found that the optimal lag maximizing the cross-mapping skill of mosquito abundance differed among climate variables (fig. 1c, left column). Significance of CCM was tested using a Mann-Kendall test for monotonic increase to examine whether increasing the number of data points used to reconstruct the system’s manifold also increased the CCM skill (Chang *et al*. 2017). In addition, a Kolmogorov-Smirnov was used to identify if this skill was significantly higher than a seasonal null model (Deyle *et al*. 2016). We defined a cross-mapping to be significant when both tests were passed. As a control, the nonsensical direction of causality was tested with the same delayed variables (fig. S3). We used the CCM implementation of the rEDM package (version 1.2.3) (Ye *et al*. 2016) to carry out all of the empirical dynamic modeling.

#### Ordinary differential equation model of larval development

We constructed a base mechanistic model for following the temperature- and density-dependent larval pupation dynamics. The model follows the ordinary differential equations (ODE):

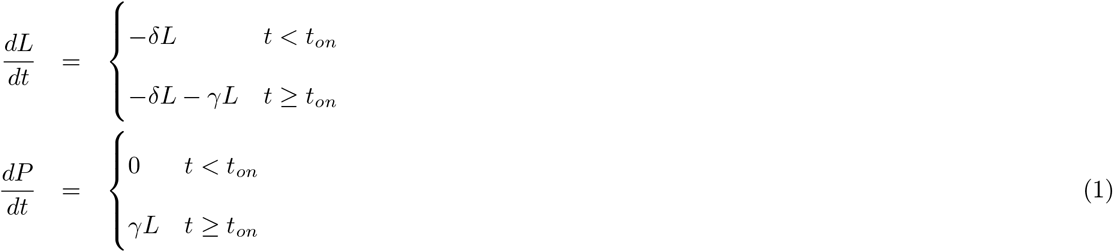

In the model, a population of larvae *L* with an initial number of *L*_0_ can die at a constant rate *δ*. Pupation starts after a time *t*_*on*_ at a transition rate *γ*, generating new pupae *P* . To understand which of these life traits are affected by the different environmental variables, we allowed them to follow distinct functional responses (table S1), producing 16 different models. We evaluated the models’ performance by using the Akaike information criterion (AIC) through fitting each model to the experimental data, which ranks each model according to the goodness of fit while adjusting for model parsimony. The best-performing model followed the equations:

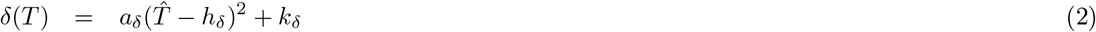

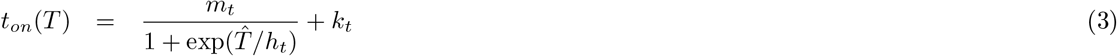

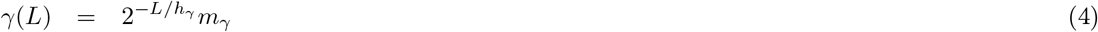

where *a*_*δ*_ is the sensitivity of death rate *δ* to temperature, *h*_*δ*_ is the temperature where the death rate is at its minimum *k*_*δ*_, *t*_*on*_ reaches its maximum *m*_*t*_ or minimum *k*_*t*_ with a decreasing rate 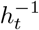, *h*_*γ*_ is the larval density at which the transition rate is half its maximum *m*_*γ*_. For the stability of the fits, we used a reparametrized expression for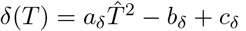, with 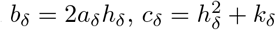and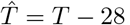 .

#### Parameter estimation

To capture the individual variability observed in the data, we used a non-linear mixed-effects framework to fit the model to the observed data of all temperature and density conditions. Here, the response of an individual pan *i* is described as

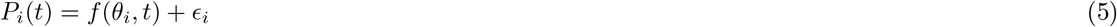

where *f* is the solution of the ODE model given by eq.1, while *θ*_*i*_ and *ϵ*_*i*_ are the coefficient vector and the residual error for pan *i*, respectively. Each element *j* of *θ*_*i*_ can be expressed as *θ*_*ij*_ = *μ*_*j*_ exp(*η*_*ij*_), where *μ*_*j*_ is the population estimate for the parameter (fixed effect) and exp(*η*_*ij*_) is the individual deviation from the population (random effect). The individual random effects and the residual error are assumed to follow independent normal distributions 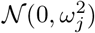 and 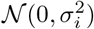, respectively.

The model parameters were estimated using Lixoft Monolix 2019R2 (Lixoft SAS 2019), using the SAEM algorithm, and the results were analyzed and visualized using R version 4.0.

#### Model validation

For validating the goodness of the fit of the best model according to AIC, we divided our data set into a training and testing set, which contained five and nine randomly selected experimental repeats, respectively. For each combination of temperature and density, we used the estimated coefficients by the training set to generate 200 simulations, obtaining a total of 3000 development curves. The individual estimates for the coefficients *θ*_*ij*_ were generated using the definition of the mixed-effects model, 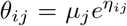, where *μ*_*j*_ is the fixed estimate for the coefficient and 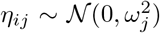 is the random effect. The biological parameters *δ, t*_*on*_, and *γ* were calculated using the definitions in eqs. 2-4, respectively. In the case of obtaining a negative value for any of the biological parameters, *θ*_*ij*_ was resampled until a positive value was obtained. Furthermore, we simulated in a similar fashion two additional experiments that were performed at 20 and 1000 Larvae/pan, both at 28°C, whic are conditions that were not used for the model fitting. To assess the goodness of the fits qualitatively, we overlayed the test set pupation curves to the 95% confidence interval of the simulations and examined how well the data and its variability were recovered by the model.

## Acknowledgements

The authors wish to thank Liane Spohr for mosquito breeding assistance. We are grateful to all members of the Vector Biology Unit for fruitful discussions and to Dr. Angelo Valleriani for helpful comments.

## Additional information

### Author Contributions

PCB, CH, AT, and EAL conceived the study. JE performed the computational modeling and data analysis. AMW performed the statistical analysis. PCB and AMW designed the laboratory experiments. AMW, PS, and CN performed the laboratory experiments. MG, AD, MD, and DS conducted the field mosquito collections and genotyping. JE, AMW, CH, PCB, and EAL contributed to writing and revising the manuscript.

### Data accessibility statement

The raw data and the code used for the analysis can be found under https://gitlab.mpcdf.mpg.de/vectorbiology/larvalthermo_mosquitodynamics.

## Supplementary Tables

**Table S1:** Akaike information criteria (AIC) of model fitting for different combinations of functional responses to temperature and larval density

| | $\delta(T, L)$ | $t_{on}(T, L)$ | $\gamma(T, L)$ | AIC |
| --- | --- | --- | --- | --- |
| 1 | $\delta$ | $t_{on}$ | $\gamma$ | -193850.0 |
| 2 | $a_\delta(\hat{T} - h_\delta)^2 + k_\delta$ | $t_{on}$ | $\gamma$ | -194251.1 |
| 3 | $a_\delta(\hat{T} - h_\delta)^2 + k_\delta$ | $-a_t\hat{T} + k_t$ | $\gamma$ | -194369.1 |
| 4 | $a_\delta(\hat{T} - h_\delta)^2 + k_\delta$ | $-a_t\hat{T} + k_t$ | $\gamma_0 e^{-L/h_\gamma}$ | -195700.3 |
| 5 | $a_\delta(\hat{T} - h_\delta)^2 + k_\delta$ | $\frac{-a_t\hat{T}}{1+\exp(-h_t L)} + k_t$ | $\gamma_0 e^{-L/h_\gamma}$ | -195668.1 |
| 6 | $a_\delta(\hat{T} - h_\delta)^2 + k_\delta$ | $\frac{m_t}{1+\exp(\hat{T}/h_t)} + k_t$ | $\gamma$ | -194633.0 |
| 7 | $a_\delta(\hat{T} - h_\delta)^2 + k_\delta$ | $\frac{m_t}{1+\exp(\hat{T}/h_t)} + k_t$ | $\gamma_0 e^{-L/h_\gamma}$ | -196384.5 |
| 8 | $a_\delta(\hat{T} - h_\delta)^2 + k_\delta$ | $\frac{m_t}{1+\exp(\hat{T}/h_t)+\exp(-L/h_{t'})} + k_t$ | $\gamma_0 e^{-L/h_\gamma}$ | -195227.2 |
| 9 | $a_\delta(\hat{T} - h_\delta)^2 + k_\delta$ | $a_t(\hat{T} - h_t)^2 + k_t$ | $\gamma_0 e^{-L/h_\gamma}$ | -195980.5 |
| 10 | $a_\delta(\hat{T} - h_\delta)^2 + k_\delta$ | $\frac{a_t(\hat{T}-h_t)^2}{1+\exp(-L/h_{t'})} + k_t$ | $\gamma_0 e^{-L/h_\gamma}$ | -195898.4 |
| 11 | $\delta_0 e^{L/h_\delta}$ | $t_{on}$ | $\gamma$ | -194632.8 |
| 12 | $\delta_0 e^{L/h_\delta}$ | $-a_t\hat{T} + k_t$ | $\gamma$ | -194685.7 |
| 13 | $\delta_0 e^{L/h_\delta}$ | $\frac{-a_t\hat{T}}{1+\exp(-h_t L)} + k_t$ | $\gamma$ | -194846.3 |
| 14 | $\delta_0 e^{L/h_\delta}$ | $-a_t\hat{T} + k_t$ | $\gamma_0 e^{-L/h_\gamma}$ | -195178.2 |
| 15 | $\delta_0 e^{L/h_\delta}$ | $\frac{-m_t\hat{T}}{1+\exp(-h_t L)} + k_t$ | $\gamma_0 e^{-L/h_\gamma}$ | -195489.3 |
| 16 | $\frac{a_\delta\hat{T}^2 L}{h_\delta} - b_\delta\hat{T} + k_\delta$ | $\frac{-a_t\hat{T}}{1+\exp(-h_t L)} + k_t$ | $\gamma_0 e^{-L/h_\gamma}$ | -195859.8 |

## Supplementary Figures

**Figure S1:**
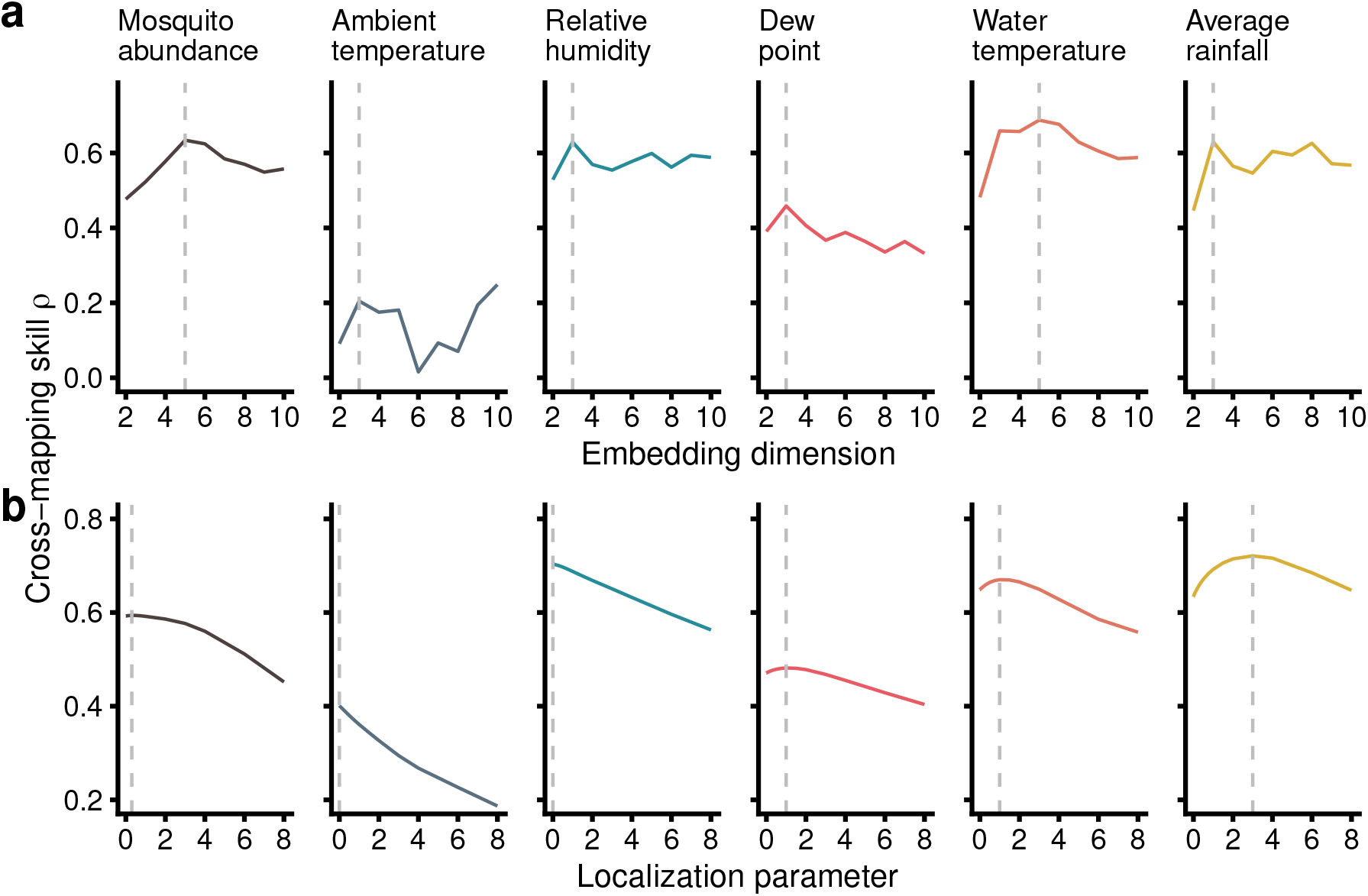
Results of the univariate state space reconstruction (SSR). **(a)** The embedding vector length was determined by selecting the *E* that maximizes the prediction skill under the range of *E* = [2, 10]. We found *E* was 5, 3, 3, 3, 5, and 3 for the adult mosquito, ambient temperature, relative humidity, dew point, water temperature, and rainfall SSRs, respectively. **(b)** The non-linearity of each variable was determined by S-map prediction varying the localization parameter *θ* = [0, 8]. Dew point, water temperature, and rainfall had a clear maximum when *θ >* 0, while the maxima for ambient temperature and relative humidity were located at *θ* = 0. The vertical dashed line indicates the maximum.

**Figure S2:**
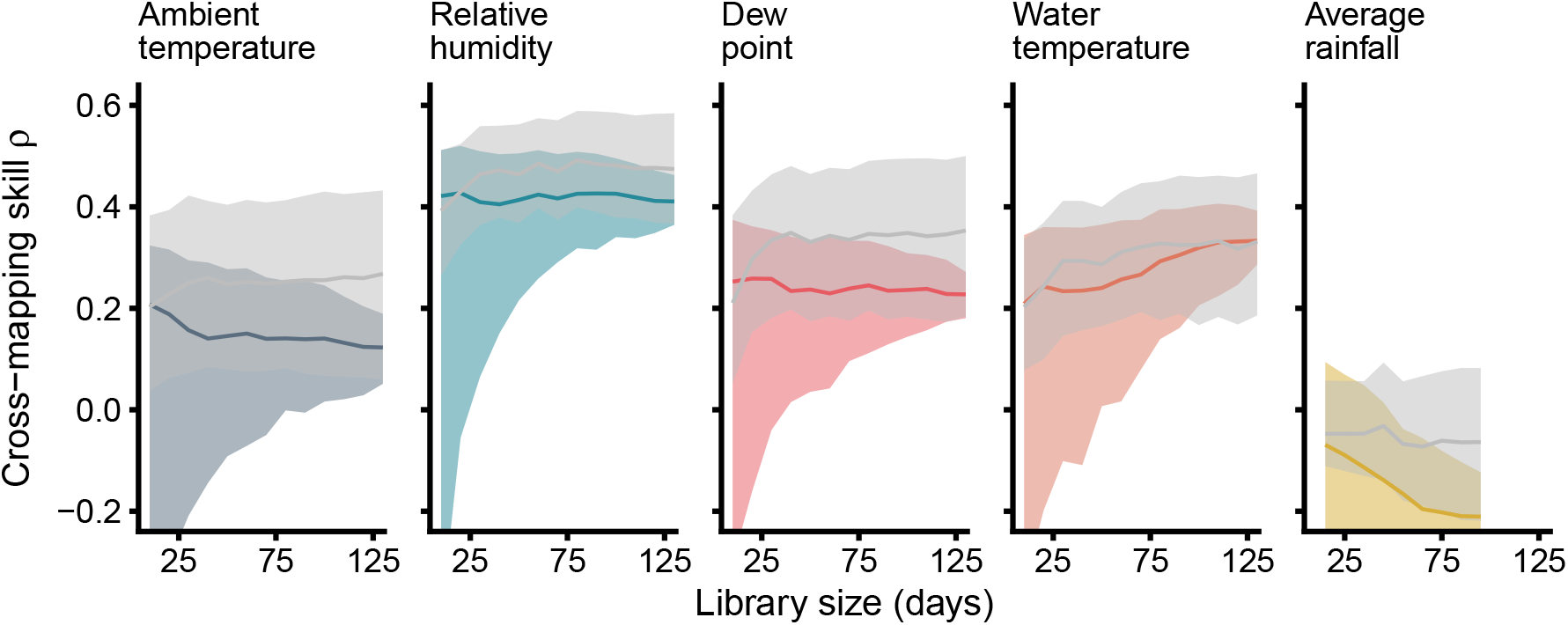
CCM with non-lagged climate variables (from left to right: ambient temperature, relative humidity, dew point, water temperature, and rainfall). This corresponds to the case where climate influences immediately the adult mosquito population. For none of the tests the cross-mapping skill was converging or different than the null model.

**Figure S3:**
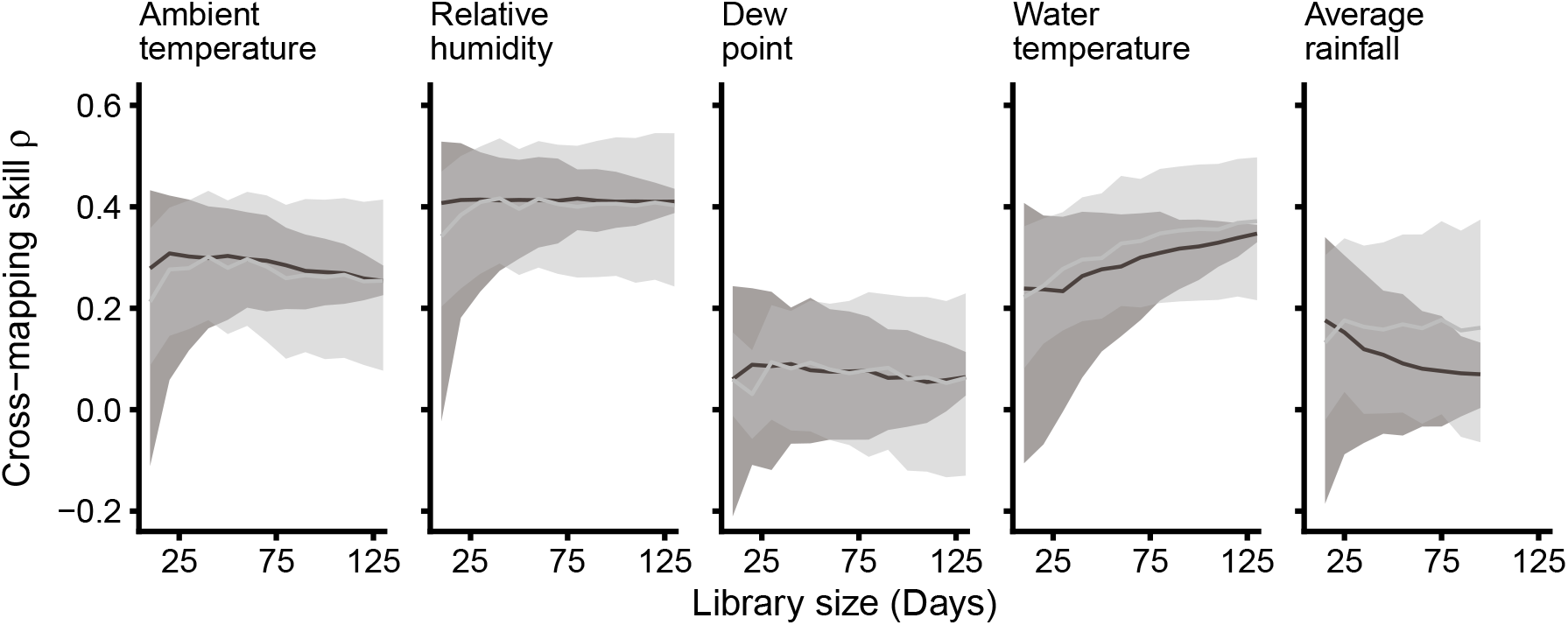
CCM control test in the non-sensical direction, where mosquitoes would be influencing the different climate variables (from left to right: ambient temperature, relative humidity, dew point, water temperature, and rainfall). For none of the tests the cross-mapping skill was converging or different than the null model.

**Figure S4:**
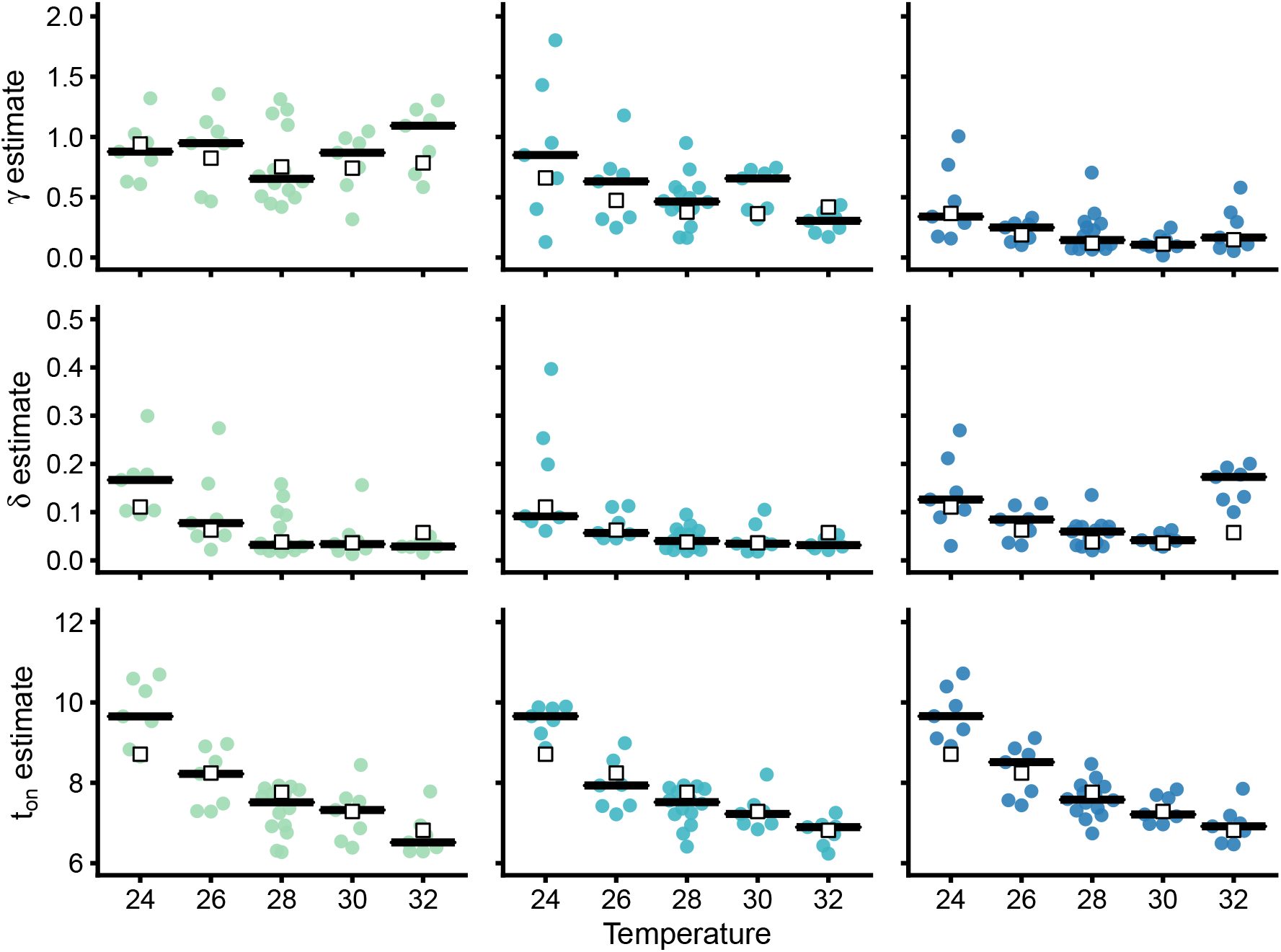
Parameter estimates from mixed-effects modeling. In general, the rate *γ* varies strongly with density, while death rate *δ* and the starting day of pupation *t*_*on*_ vary strongly with temperature. Individual estimates for the different experimental repeats are represented by colored points. The median of the individual estimates for each condition (black line) correlates strongly with the population estimates (fixed effects, blank square).

**Figure S5:**
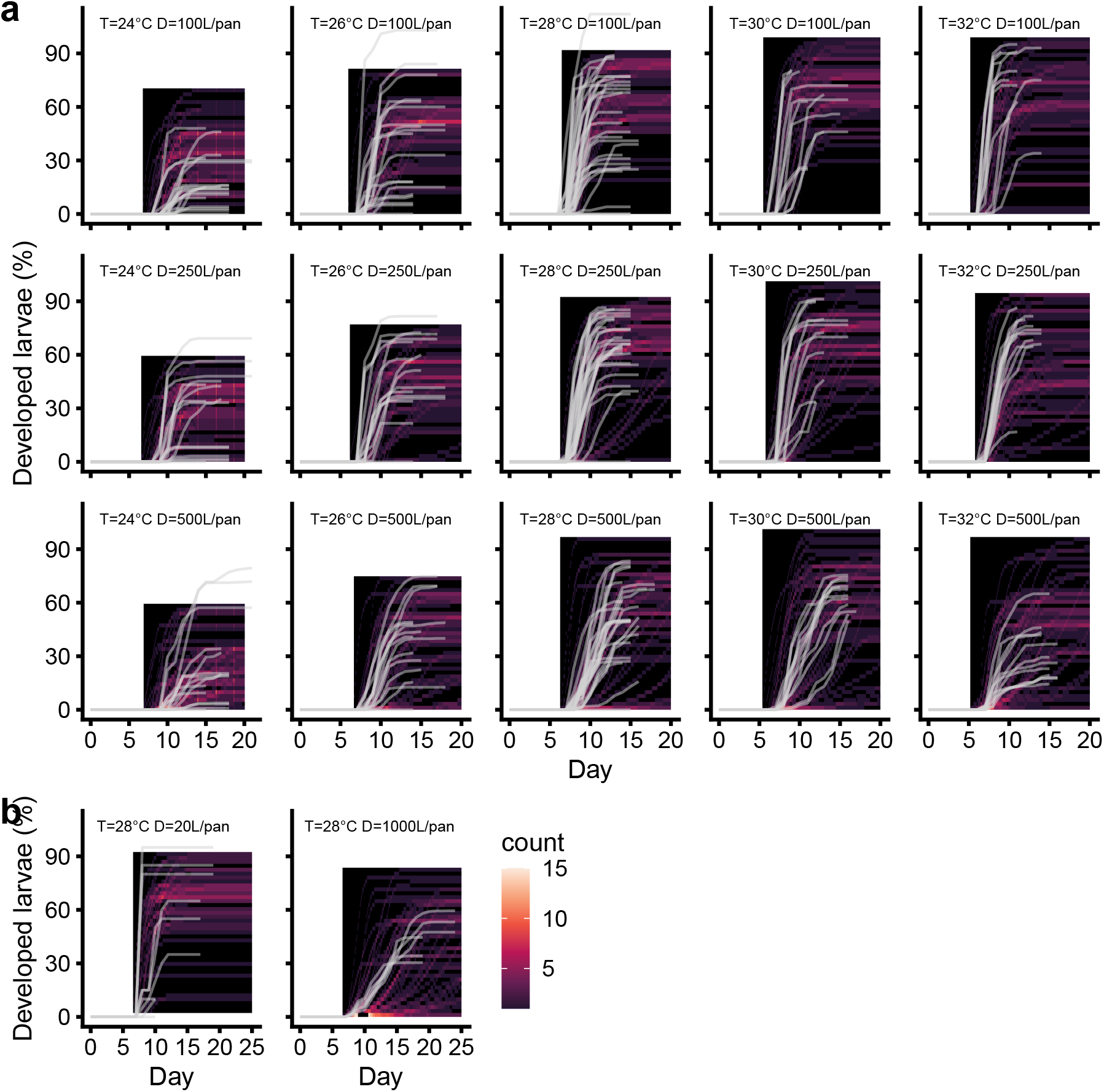
Qualitative validation of mixed-effects model for larval development. **(a)** Black lines represent test data from nine experimental repeats, while the heat-map colors represent the overall behavior of 50 simulated development curves. The parameters were obtained by fitting the remaining five repeats. In general, all pupation curves follow the general trend of the simulations, meaning that the model captures well the dynamics of larval development. **(b)** Out-of-sample validation, gray lines represent out-of-sample data from three experimental repeats performed at 28°C with 20 or 1000 larvae per pan. The parameters were obtained by fitting all repeats from the conditions in the larval development experiment. Overall, the model captures well the behavior of low and high larval densities, but overestimates the slow development on high densities.

